# RTFED, an open-source versatile tool for home-cage monitoring of behaviour and fibre photometry recording in mice

**DOI:** 10.1101/2025.07.24.666497

**Authors:** Hamid Taghipourbibalan, James Edgar McCutcheon

## Abstract

**Background:** Conventional approaches for studying feeding and reward-driven behaviours require frequent animal handling or relocation of animals to specialized chambers, inducing stress, confounding behavioural outcomes, and limiting continuous (24/7) data collection. In recent years, the Feeding Experimentation Device (FED3) has emerged as a major advance, offering programmable modes of operation, affordable costs, and flexibility for investigating a range of feeding and operant behaviours. However, certain limitations prevent researchers from fully harnessing the FED3’s capabilities in a user-friendly manner.

**New method:** Here, we present the Realtime and Remote FED3 (RTFED) developed for continuous and online home-cage monitoring of mice, video recording behaviours and fibre photometry recording.

**Results:** Validation experiments confirm RTFED integrates well with FED3 to log and transmit behavioural events in real-time. It also incorporates event-triggered video capture through USB cameras, providing additional observational depth. Moreover, RTFED handles TTL signals to the fibre photometry system allowing precise behaviour-neural synchronization.

**Comparison with existing methods:** A key strength of RTFED is its easily customizable architecture, enabling researchers to tailor both software and hardware configurations to meet specific experimental objectives. This flexibility, together with features such as remote data logging and email notifications that allow timely adjustments and animal welfare monitoring based on behavioural observations, substantially reduces animal disturbance and researcher intervention and labour.

**Conclusions:** By offering a cost-effective and modifiable alternative to proprietary commercial solutions, RTFED broadens accessibility, heightens reproducibility, and deepens investigations into feeding and reward-driven behaviours in home-cage settings, ultimately improving the quality and translational relevance of behavioural research.

**Highlights:**

- We enhanced the utility of FED3 with a graphical user interface called RTFED
- RTFED allows real-time and remote monitoring of mouse feeding and operant behaviour
- It sends notifications in case of device failures or when flagged behaviours happen
- It records event-triggered videos of defined behaviour
- RTFED relays sub-millisecond TTL outputs for fiber photometry recording

**Graphical abstract:** 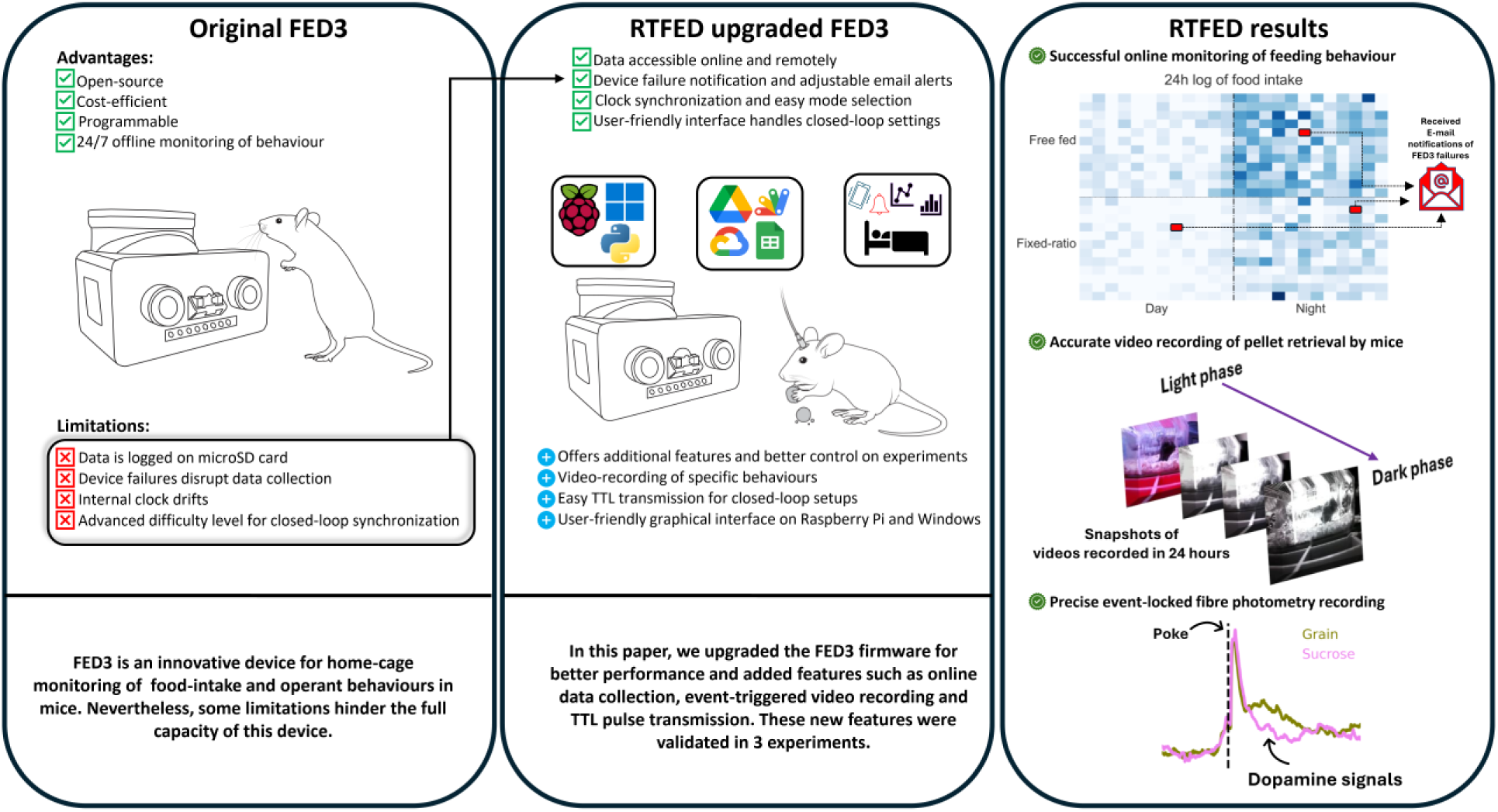

## 1. Introduction

Quantifying rodent behaviour in consistent and ethologically relevant ways remains a central challenge in neuroscience (Levitis et al., 2009), particularly given the need for accurate monitoring of behaviours such as feeding patterns and operant responses, which are critical for uncovering the neural circuits and physiological mechanisms underlying homeostasis, metabolism, and reward-driven processes (Berthoud & Münzberg, 2011; Morrison et al., 2012). Traditional approaches often relied on manual measurement of parameters and brief tests capturing short snapshots of behaviour in specialized environments, where researchers physically monitored animals in controlled conditions such as operant chambers or open-field arenas, and logged behaviours like lever presses, reward retrieval, or locomotor activity (Skinner, 1938; Ellenbroek & Youn, 2016). While such methods have yielded fundamental insights into understanding behaviour, they also introduced significant limitations. For example, rodents tested outside their home-cage experience novelty-induced stress and anxiety to the artificial context, which can disrupt naturalistic behaviour. In addition, the presence or intervention of human observers can add variability and bias to behavioural measurements (Ali & Kravitz, 2018; Francois et al., 2022; Snyder et al., 2024). Furthermore, such approaches often demand substantial time and effort by the experiment (Kahnau et al., 2023).

The advent and development of modern automated home-cage monitoring systems initiated a paradigm shift, allowing continuous observation of animals in their familiar environments over extended time periods via the aid of electronic sensors, modules and video tracking, thus minimizing unwanted and uncontrolled factors in addition to enhancing ecological validity of experiments (Richardson, 2015; Steele et al., 2007). Since then, robust behavioural monitoring in home-cage environments has become indispensable for studying naturalistic behaviours such as food intake, reward-seeking, circadian rhythms, and decision-making along with measurements of several biological factors in rodents (Goulding et al., 2008; Kahnau et al., 2023). Moreover, in recent years these innovative technologies incorporate artificial intelligence and machine learning models to unravel aspects of behaviour that remained unclear and understudied for decades (Isik & Unal, 2023; Mathis et al., 2018; Pereira et al., 2022)

Commercially available home-cage monitoring systems, while effective, are often prohibitively expensive and closed-source. This limits their adaptability and transparency, and might discourage smaller laboratories or institutions with limited funding from pursuing cutting-edge research, thereby constraining innovation in behavioural science (Mingrone et al., 2020). These proprietary systems can be difficult to customize or freely integrate with other experimental setups, restricting their utility in complex, multi-modal studies that require the combination of behavioural data with other physiological measurements (Singh et al., 2019). This situation has driven the development of open-source alternatives made by researchers for researchers, which prioritize affordability, transparency, and adaptability (e.g., (Benedict & Cudmore, 2023) , (Godynyuk et al., 2019), (Virag et al., 2021) and OpenBehavior Project, n.d.). One of the tools developed in response to this condition was the Feeding Experimentation Device 3 (FED3) that has significantly transformed the study of food intake and operant behaviour in rodents by providing an open-source, low-cost, and customizable alternative to commercial systems. This device was developed as a standalone, battery-powered, 3D printable and DIY tool allowing researchers to measure and manipulate feeding and reward-seeking behaviour in freely moving rodents while automatically tracking parameters such as pellet retrieval and nose pokes (Matikainen-Ankney et al., 2021; Nguyen et al., 2017). However, despite the capabilities of FED3, a few limitations have hindered the complete utility of this device. For example, FED3 stores data on a micro-SD card making these data inaccessible during the experiment and thus limiting concurrent analysis. Moreover, occasionally the devices can become clogged resulting in “jamming” events and preventing delivery of pellets, which causes data loss or animal welfare issues. In addition, the internal clock of FED3 drifts over time such that synchronization across multiple devices is difficult and inaccurate. Finally, FED3 is in theory capable of synchronizing with other devices (e.g., neural recording hardware) via a BNC/minijack port (Krynitsky et al., 2020; Matikainen-Ankney et al., 2021). However, implementation of such synchronization requires a level of technical knowledge and customization that may be beyond users with a limited coding and electronics background.

Here, we introduce the Real-time and Remote FED3 (RTFED) system, a software package requiring only minor hardware modifications to level-up the capabilities of FED3 devices in a highly user-friendly manner. This system runs on both Windows OS and Raspberry Pi, facilitates real-time and remote data collection, event-triggered video recording of behaviour, and transmission of Transistor-Transistor Logic (TTL) pulses to external devices (here, Tucker Davis Technologies fibre photometry systems). Designed as a user-friendly upgrade to existing FED3 devices, RTFED allows researchers, even those with limited experience in programming and electronics, to easily implement customized experimental protocols and enhance their research. Here we conducted three sets of experiments to validate the functionality and reliability of the system and demonstrate that the RTFED is a versatile and suitable tool for deployment in both home-cage environments and controlled experimental setups.

## 2. Materials and methods

### 2.1 Animals and procedures

To verify the functionality of the system, a total of 12 adult male C57BL/J6 mice (6-8 weeks old, weighing 21-30 g) were purchased from Janvier Labs (France), acclimatized for 5 days in a temperature (22 ± 0.5 °C) and humidity (56 ± 2%) controlled room, under a 12h light/dark cycle (lights on 12:00 AM, lights off 12:00 PM). Subsequently, mice were moved to another room with similar environmental conditions where the experiments were conducted. Two animals per cage were contact-housed in modified conventional cages (1284L EUROSTANDARD TYPE II L, TECNIPLAST, Italy) with dimensions of 365 x 207 x 140 mm and floor area of 530 cm^2^ separated in half by a perforated divider that allowed visual, olfactory and minor physical communication but no major physical contact. Each cage was filled with ∼1 kg of wood shaving as cage bedding and each mouse was provided with 3.65 g cotton balls as nesting material. Water and food (regular chow pellets; ssniff Rat/Mouse – Maintenance diet) were available ad libitum otherwise mentioned. All animal care and experimentation were in compliance with the EU directive 2010/63/EU for animal experiments and were approved by the Norwegian Food Safety Authority (FOTS #30889).

### 2.2 The overview of the RTFED system

The RTFED setup (Fig. 1) is a Python-based software designed to monitor and log data from FED3 devices directly to a Google spreadsheet. It enables real-time monitoring of feeding and operant behaviour in mice, making it particularly suitable for behavioural neuroscience experiments conducted in home-cage settings. Moreover, when incorporated with a Raspberry Pi, RTFED is also capable of coupling behavioural events of interest (e.g., pellet intake) with video recording. Additionally, RTFED enables users to relay behavioural events as TTL signals to external devices allowing, for example, fibre photometry recordings to be synchronized with behavioural events within the home-cage. Recommended items needed to run RTFED on Windows and Raspberry Pi are listed in Table 1.

**Fig. 1.**
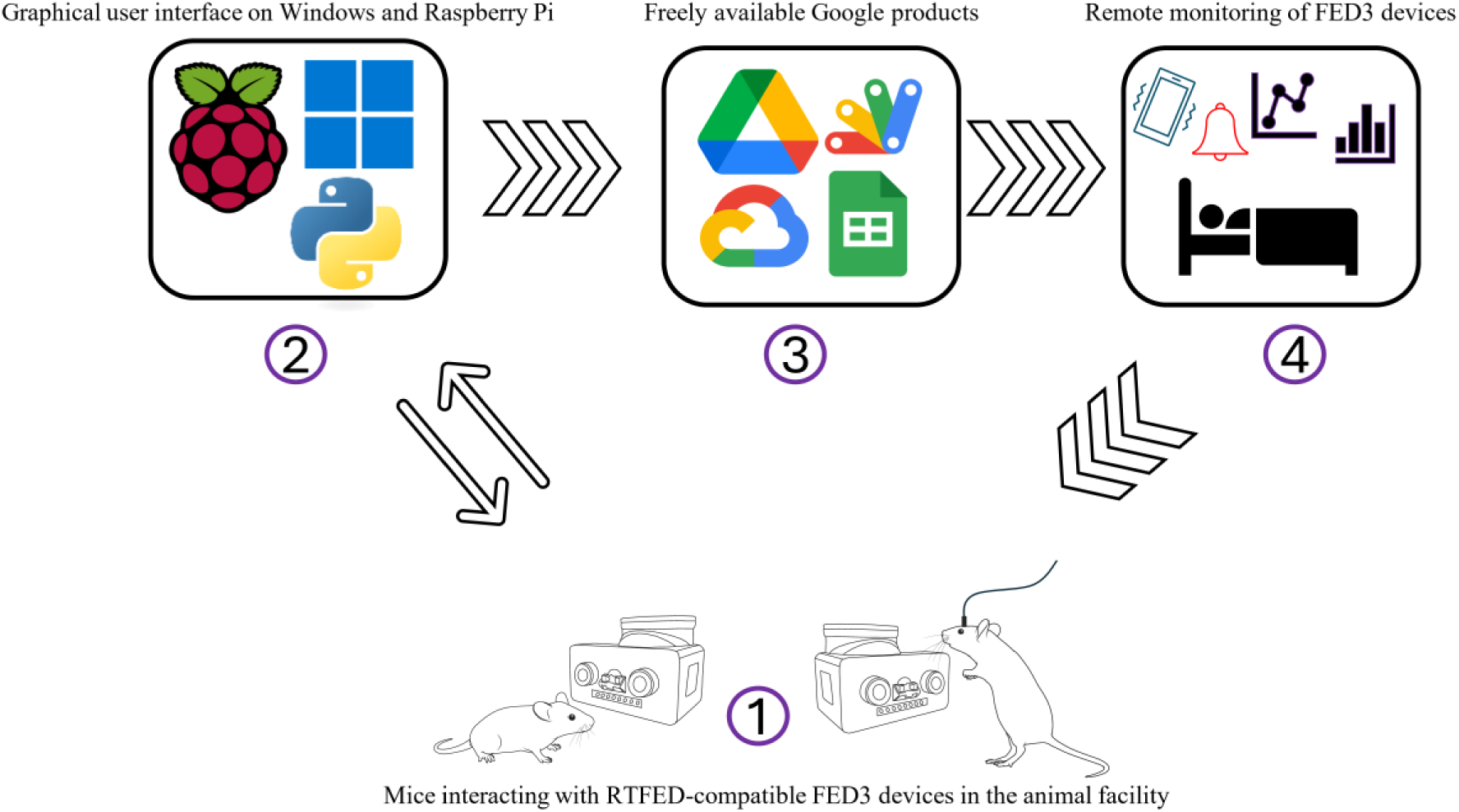
When running on Windows OS, the RTFED software monitors interactions with FED3 devices online and remotely and data are transmitted to a Google spreadsheet, which is accessible to the user on all compatible platforms. By using Google Apps Scripts, it is possible to define customized email notifications according to desired behavioural outputs or further process and plot the data. The same features are available on a Raspberry Pi OS, but additionally it is possible to have event-triggered video recordings of behaviour or TTL pulse transmission.

**Table 1.**
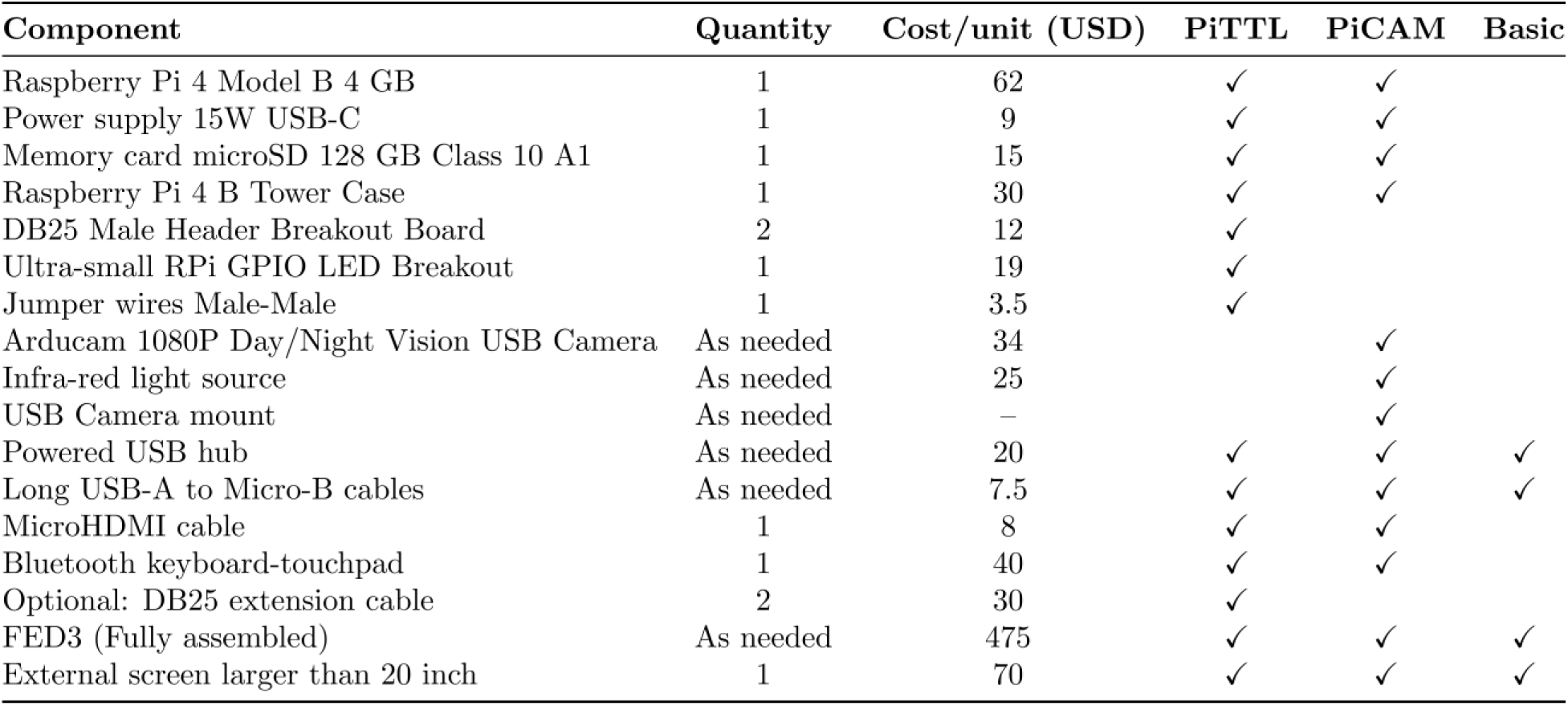
The list of items needed to setup full-featured RTFED including the Basic RTFED, PiCAM and PiTTL modules. The cost per unit is approximately estimated based on geographical location. Suggested links to the items can be found on our GitHub repository.

| Component | Quantity | Cost/unit (USD) | PiTTL | PiCAM | Basic |
| --- | --- | --- | --- | --- | --- |
| Raspberry Pi 4 Model B 4 GB | 1 | 62 | ✓ | ✓ |  |
| Power supply 15W USB-C | 1 | 9 | ✓ | ✓ |  |
| Memory card microSD 128 GB Class 10 A1 | 1 | 15 | ✓ | ✓ |  |
| Raspberry Pi 4 B Tower Case | 1 | 30 | ✓ | ✓ |  |
| DB25 Male Header Breakout Board | 2 | 12 | ✓ |  |  |
| Ultra-small RPi GPIO LED Breakout | 1 | 19 | ✓ |  |  |
| Jumper wires Male-Male | 1 | 3.5 | ✓ |  |  |
| Arducam 1080P Day/Night Vision USB Camera | As needed | 34 |  | ✓ |  |
| Infra-red light source | As needed | 25 |  | ✓ |  |
| USB Camera mount | As needed | – |  | ✓ |  |
| Powered USB hub | As needed | 20 | ✓ | ✓ | ✓ |
| Long USB-A to Micro-B cables | As needed | 7.5 | ✓ | ✓ | ✓ |
| MicroHDMI cable | 1 | 8 | ✓ | ✓ |  |
| Bluetooth keyboard-touchpad | 1 | 40 | ✓ | ✓ |  |
| Optional: DB25 extension cable | 2 | 30 | ✓ |  |  |
| FED3 (Fully assembled) | As needed | 475 | ✓ | ✓ | ✓ |
| External screen larger than 20 inch | 1 | 70 | ✓ | ✓ | ✓ |

### 2.3 RTFED library as an upgrade to the original FED3 firmware

The initial step in the process of upgrading a regular FED3 unit to an RTFED-compatible unit is to update the existing FED3 library to handle commands to and from the host computer. These enhancements ensure FED3 firmware can communicate with the RTFED GUI, and that the user can utilize all the features provided by RTFED. The detailed modifications made to the firmware can be found in the Supplementary file, sections 2.3.S1 to S5, but a brief summary of the differences under RTFED firmware is reported on Table 2.

**Table 2.** A comparison between features available on the original FED3 compared with the same devices flashed with RTFED library. The tweaks in the firmware open up lots of possibilities beyond the initial capabilities of FED3.

| Feature | FED3 | RTFED |
| --- | --- | --- |
| Data logging | Local SD card; post-hoc retrieval | Real-time stream to PC/Raspberry Pi; optional cloud/CSV backup |
| Real-time view | No (stand-alone logger) | Yes (live event stream) |
| Sync / I/O | Limited / user add-ons | Built-in TTL for photometry / video recording / closed loop |
| Parallel devices | Per-unit stand-alone | 10–20+ devices in parallel via USB hubs |
| Power | Battery powered | USB powered (PC or Raspberry Pi) |
| Data export | CSV on SD | CSV on SD and locally + live Google Sheets |
| Best use case | Offline home-cage feeding or operant training | Multi-device, real-time, closed-loop / photometry-synced experiments |

### 2.4 Google console project

After flashing a FED3 device with the RTFED library, to enable remote data logging and automated notifications, the RTFED system should be integrated with a Google Cloud Console project. Within this project, a service account is created and granted permission to access Google Sheets and Google Drive via their respective APIs. A JSON credential file generated from the service account is used by the RTFED software to authenticate and append data to a designated Google spreadsheet in real-time. The spreadsheet is shared with the service account to allow read/write access. Additionally, a Google Apps Script is embedded within the spreadsheet to monitor incoming data for desired events (e.g., number of pellets taken or FED3 failures) and trigger email alerts automatically if predefined conditions are met. This cloud integration allows the RTFED system to operate remotely, providing real-time behavioural monitoring and timely experimenter notifications without manual oversight. The full process of setting up the project on Google console is described on our GitHub page.

### 2.5 RTFED (Basic) on Windows and Raspberry Pi

The RTFED GUI (Supplementary file, Fig. S1 A-B) is a standalone Python software designed for real-time behavioural data logging from multiple FED3 devices (Fig. 2). The GUI works as a dashboard to send commands to FED3 units, store data in structured folders and transmit the logged data directly to Google spreadsheets where the users can process it further online and remotely. By using Apps Script, they can interact with the spreadsheet, e.g., setting an alarm to notify them about certain events.

**Fig. 2.**
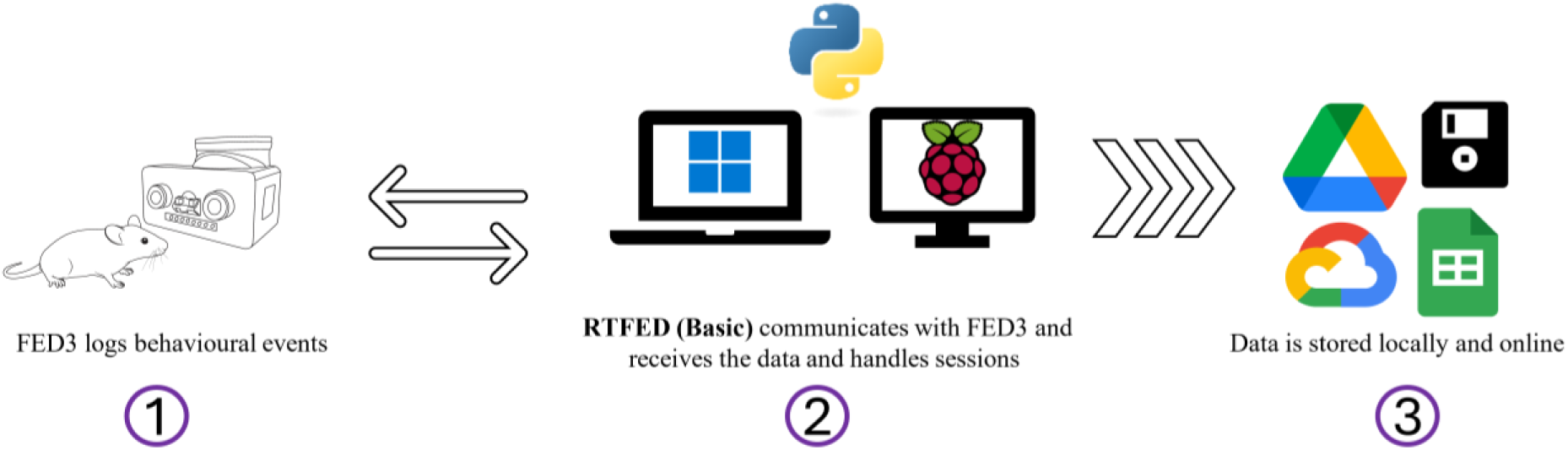
A schematic representation of RTFED (Basic). The GUI provides an interface to command-and-control FED3 units, create structured experiment sessions, and transmit the data from FED3 units to Google spreadsheets. As soon as the data is online, the user can inspect it. At the end of the session, the data is stored locally.

The RTFED software also handles interruptions in the internet connection by caching the data and transmitting online after some seconds. It also enables users to disconnect and reconnect new devices while an experiment is ongoing, which is helpful in case of device failures or if new devices needed to be introduced to the system without stopping the logging process. The system can handle as many FED3s as needed, as long as a powered USB hub is used to connect all the extra FED3s. For a video tutorial, please watch Supplementary video V1.

### 2.6 RTFED (PiCAM)

The camera-enabled RTFED which runs on a Raspberry Pi board (Supplementary file, Fig. S2. A) and is called RTFED (PiCAM) is a separate GUI (Supplementary file, Fig. S2 B) that allows users to video record certain behaviours in addition to the basic features of RTFED (Fig. 3). Here the user can select the event that triggers video recording including Pellet delivery, Left/Right pokes, or all the events together. Up to 20 USB cameras are supported. Any USB camera can be used, while we recommend using the cost-efficient, infra-red capable camera listed in Table 1. This camera integrates easily with RTFED (PiCAM) and with a customized 3D printed frame adjusts well to different types of cage racks (Supplementary file, Fig. S2 C).

**Fig. 3.**
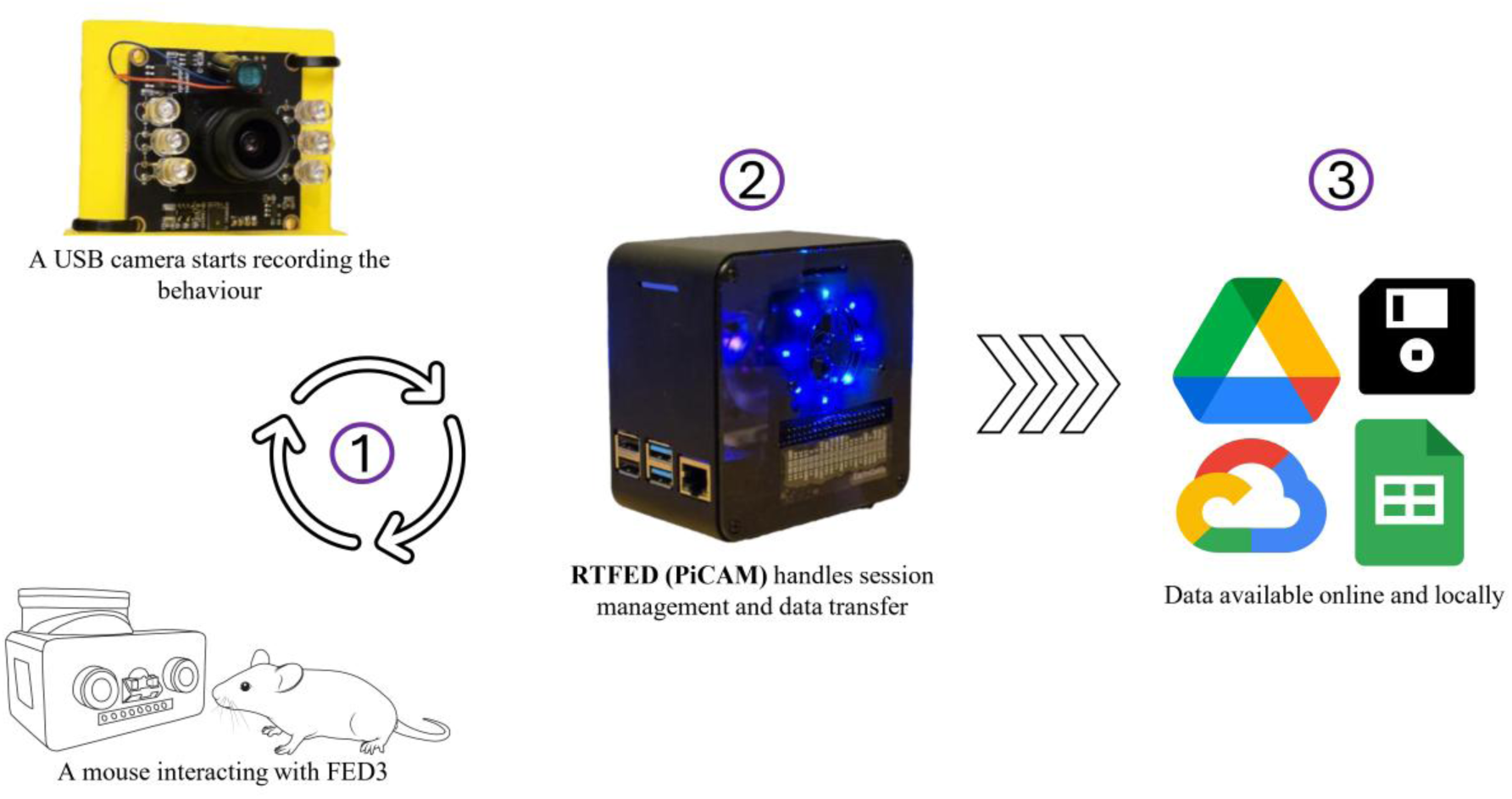
The PiCAM setup incorporates USB cameras with FED3 units. In addition to all features of RTFED (Basic), this GUI also records videos of user-defined behaviours such as pellet retrieval and nose pokes and saves the videos locally on the Raspberry Pi. The behavioural data is also transmitted online.

The RTFED (PiCAM) records 30 s videos when the selected event is logged on FED3 and in cases the event occurs again within the 30 s window, the recording is extended for another 30 s. Videos are saved once this 30 s window is reached without any further events.

### 2.7 RTFED (PiTTL)

RTFED on Raspberry Pi can also be used as a relay station in closed-loop setups to transmit TTL pulses initiated by events triggered on FED3 devices (Fig. 4, and Supplementary video V2). To accomplish this, the user only needs to connect the FED3 units to the Raspberry Pi with pins connected to a DB25 cable breakout block according to the Supplementary file Fig. S3. Each Raspberry Pi 4B can handle up to 8 FED3s at a time. This number is limited by the GPIO pins available on the Raspberry Pi.

**Fig. 4.**
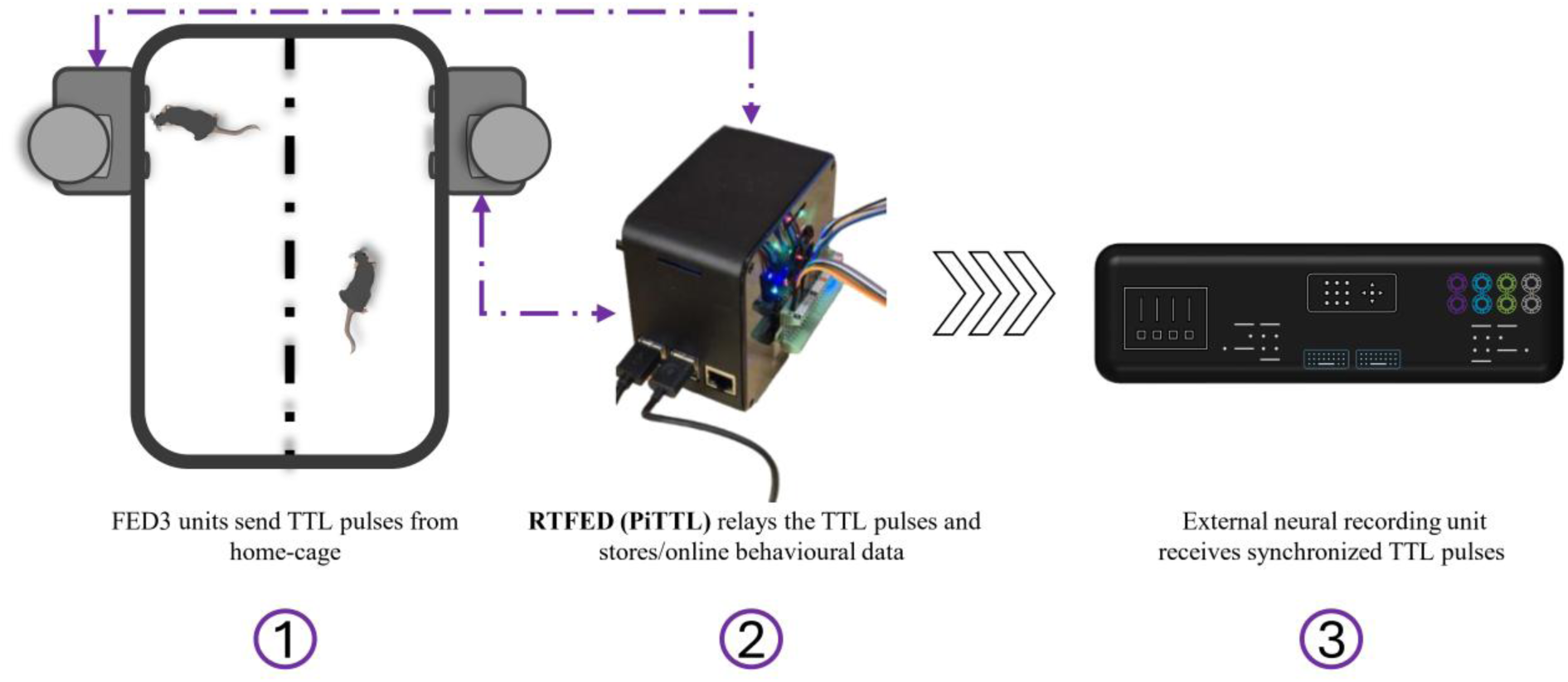
The PiTTL setup allows users to record photometry signals coupled with behavioural events on FED3 units. Up to 8 FED3 units can be handled simultaneously. Mice are contact-housed in standard cages, separated by a Plexiglas divider, the PiTTL software handles recording sessions, relays behavioural events to the neural recording systems (in this case the TDT fibre photometry processor) and stores the behavioural data locally. Optionally, the users can view the behavioural data online.

The software structure of the GUI (Supplementary file, Fig. S4) is hardwired for 8 ports and is designed to send TTL pulses (adjustable duration of 100 milliseconds) for Left/Right pokes and Pellet deliveries (Pellet Onset). The same pins handle TTL signals for LeftWithPellet and RightWithPellet events along with retrieval of the pellets (Pellet Offset). This setting means that 3 pins of the Raspberry Pi are dedicated to each FED3 unit.

### 2.8 Validation experiments

To test the functionality and feasibility of the RTFED system, we conducted three experiments. In *Experiment 1* we examined the basic functionality of RTFED to monitor food intake of mice earning 20 mg grain pellets from FED3 devices on Free Feeding mode (FF) where a food pellet is available as soon as the previous one is taken from the pellet well, and Fixed-Ratio 1 mode (FR1) where the mice have to make a single left poke to receive a pellet. Twelve male mice were contact housed 2 per cage, as described in section 2.1 and each mouse had access to its individual FED3 unit. The mice were first trained to receive 20 mg grain pellets from FED3 devices for 3 days, after training the FED3 units were connected to RTFED (Basic) running on a laptop with Windows 11 connected to an external 27 inch monitor, data was logged for ∼48 h; the first ∼24 h were dedicated to monitor food intake on FF mode and the second ∼24 h was to monitor food intake on FR1 mode.

In *Experiment 2*, we tested the reliability of the PiCAM system. The same mice from *Experiment 1* were used in this experiment and their food intake behaviour was monitored on FF mode while pellet retrieval events were used to trigger video recordings. Two mice per day were recorded. Thus, we used two FED3 units each with its own USB camera connected to the Raspberry Pi. Mice had access to regular chow in addition to grain pellets dispensed from FED3 and were free to either collect food from FED3 or from the standard food hopper in the cage.

Lastly, in *Experiment 3* to confirm the RTFED (PiTTL) can be successfully integrated with fibre photometry systems, we expressed the fluorescent dopamine sensor, *GRAB_DA_*, in the NAc core to detect dopamine release associated with behavioural events. Surgical methods were similar as reported in (Volcko et al., 2025). Briefly, under deep surgical anaesthesia (1.5-2% isoflurane suspended in air), mice were placed in a stereotaxic frame (Kopf, Tujunga, CA, USA), a small hole was drilled at the coordinates of NAc core (AP +1.5, ML +1.0, relative to Bregma and DV -4.65, relative to a flat and levelled skull) and injected with 500 nL of AAV1-hsyn-GRAB_DA2m into the NAc core with a rate of 100 nL/minute using a Nanofil syringe (33G blunt needle) and UMP3 WPI microsyringe pump (USA) and had a fibre optic cannula implanted at DV -4.55. Each mouse was given subcutaneous injections of buprenorphine (0.1 mg/kg; Temgesic) and meloxicam (5 mg/kg; Metacam) to provide systemic analgesia, as well as bupivacaine (1 mg/kg; Marcain) as a local anaesthetic. After the surgery all the mice had a week of careful follow up with daily health check and received meloxicam in the first 2 days postoperatively.

After two weeks of recovery from surgery and time for expression of the virus, the mice were maintained on normal diet (containing 20% Casein as the source of protein) for one week and subsequently recorded during the dark phase within their home-cages using the TDT RZ10X system in a 1-hour session with FED3s on FR1 mode, while receiving grain pellets (Bio-Serv, F0163). 1 day later, the same mice were recorded in their home-cages with FED3 on FR1 mode receiving sucrose pellets (Bio-Serv, F07595), (Supplementary file, Fig. S5 A-B and supplementary video V3).

### 2.9 Code availability

Code for all projects is hosted in the GitHub repository with full documentation and equipment needed: https://github.com/mccutcheonlab/FED_RT. The repository contains two branches */main* and */RTFEDPi* that can be access from the home page and also includes a link to Raspberry Pi image file that can be used to clone the system(s) used in the paper. The users can access the GitHub page to follow instructions for setting up RTFED in their labs.

## 3. Results

The three experiments conducted to validate the functionality of the RTFED GUI package allowed us to verify the setup and demonstrated its reliability.

### 3.1 Experiment 1: Home-cage monitoring of Free Feeding and FR1 modes with RTFED (Basic)

We used RTFED to log data from the two days of our experiment using FF and FR1 modes (Fig. 5**A**). As expected, and shown in other studies (Barrett et al., 2025), mice collected more pellets during FF mode than in FR1 mode. Analysis of FED3 data with respect to time of day shows that mice retrieve more food pellets during the first half of the dark phase (Fig. 5**B**). Moreover, the analysis of learning performance on FR1 mode (Fig. 5**C**) shows that mice rapidly learn to poke in the active port more than the inactive port once exposed to FR1 mode. This behaviour is in accordance with other studies carried out with FED3 (Matikainen-Ankney et al., 2021).

**Fig. 5.**
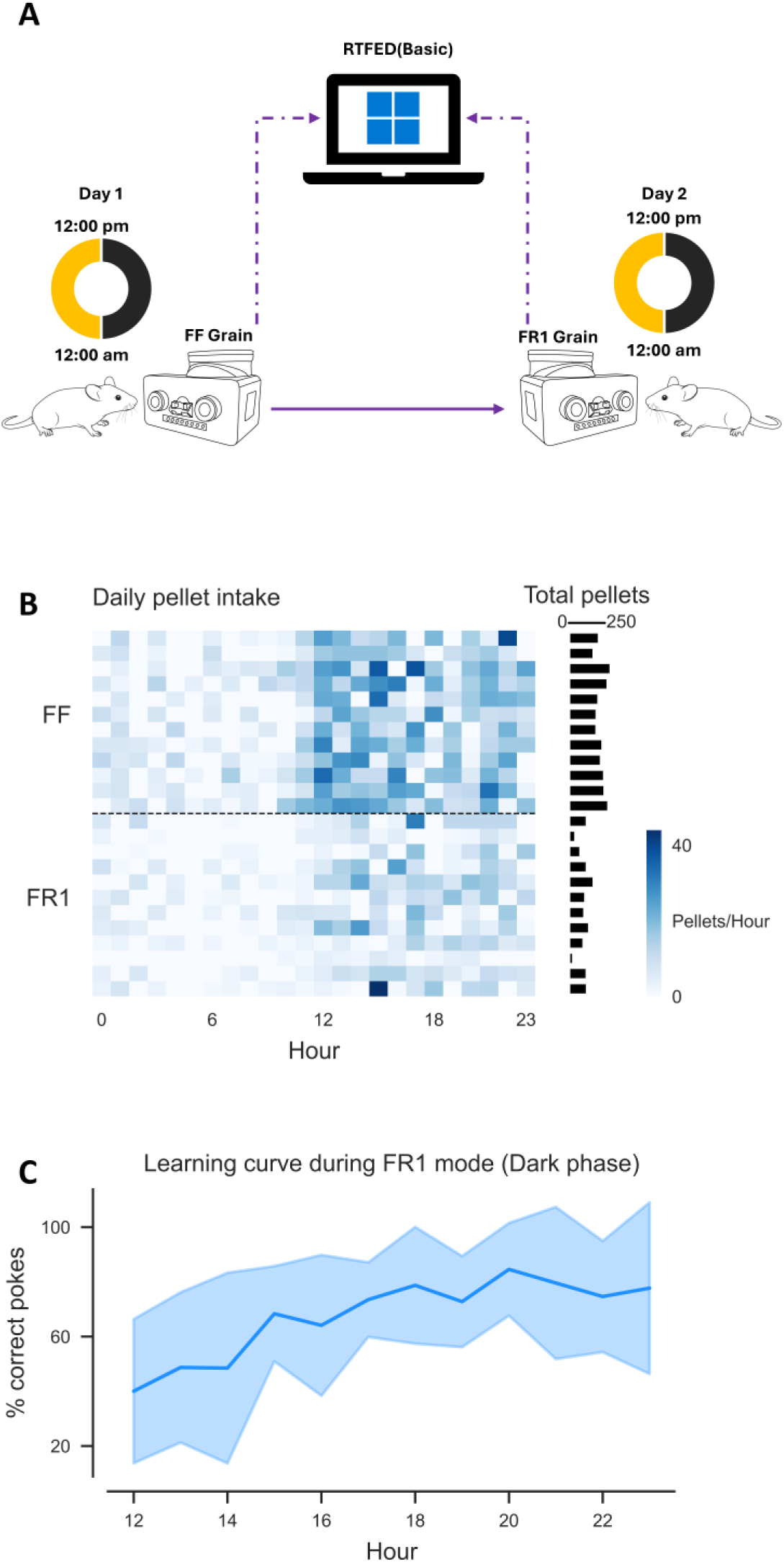
Validation experiment of home-cage monitoring with RTFED (Basic). (A) A representation of the experimental paradigm, each mouse had access to its own FED3 unit connected to the host computer, ∼24 hours on FF mode and ∼24 hours on FR1 mode obtaining grain pellets. (B) The heatmap (left panel) shows number of pellets taken per hour across a 24 h period with rows showing data from individual mice. Feeding activity follows a circadian pattern, 0-12 light phase and 12-23 dark phase. The horizontal bar plot (right panel), with each bar corresponding to a row in the heatmap, highlights reduced pellet retrievals when mice are on FR1 mode, relative to FF mode. (C) Poke efficiency during the FR1 session reflecting learning performance of mice during the dark phase. Thick line is mean and shaded area is SEM.

The RTFED (Basic) software worked without any problem on a Windows computer. Data were successfully logged from twelve FED3 units and transmitted to the Google spreadsheet. We only had 1 case of jamming which was successfully reported by the RTFED software resulting in an email to the responsible scientist allowing the issue to be rapidly resolved. This experiment confirmed that RTFED - and the data logged by it - can be reliably used for studying food intake and other relevant behaviours within the home-cage.

### 3.2 Experiment 2: A surveillance of pellet retrieval using RTFED (PiCAM)

In this experiment, pellet-triggered video recording was implemented using the RTFED (PiCAM) system during a ∼24h session on FF mode (Fig. 6**A**-**B**), where each pellet delivery initiated a 30 s video capture. If additional pellets were collected within the 30 s window, the recording was extended accordingly (Supplementary video, V4). Thus, temporally clustered pellet events resulted in a single continuous video. To verify the reliability of the PiCAM system in recording pellet-triggered videos, we calculated the number of temporally clustered pellet events per mouse, using a 30-second inter-pellet interval as a cutoff. Thus, these clusters represent the times when we expected video recording to be triggered. The number of detected clusters exactly correlated with the actual number of video files recorded for each mouse (Fig. 6**C**). As such, all video counts perfectly matched the expected values derived from the feeding logs, confirming high temporal precision and reliability of the system.

**Fig. 6.**
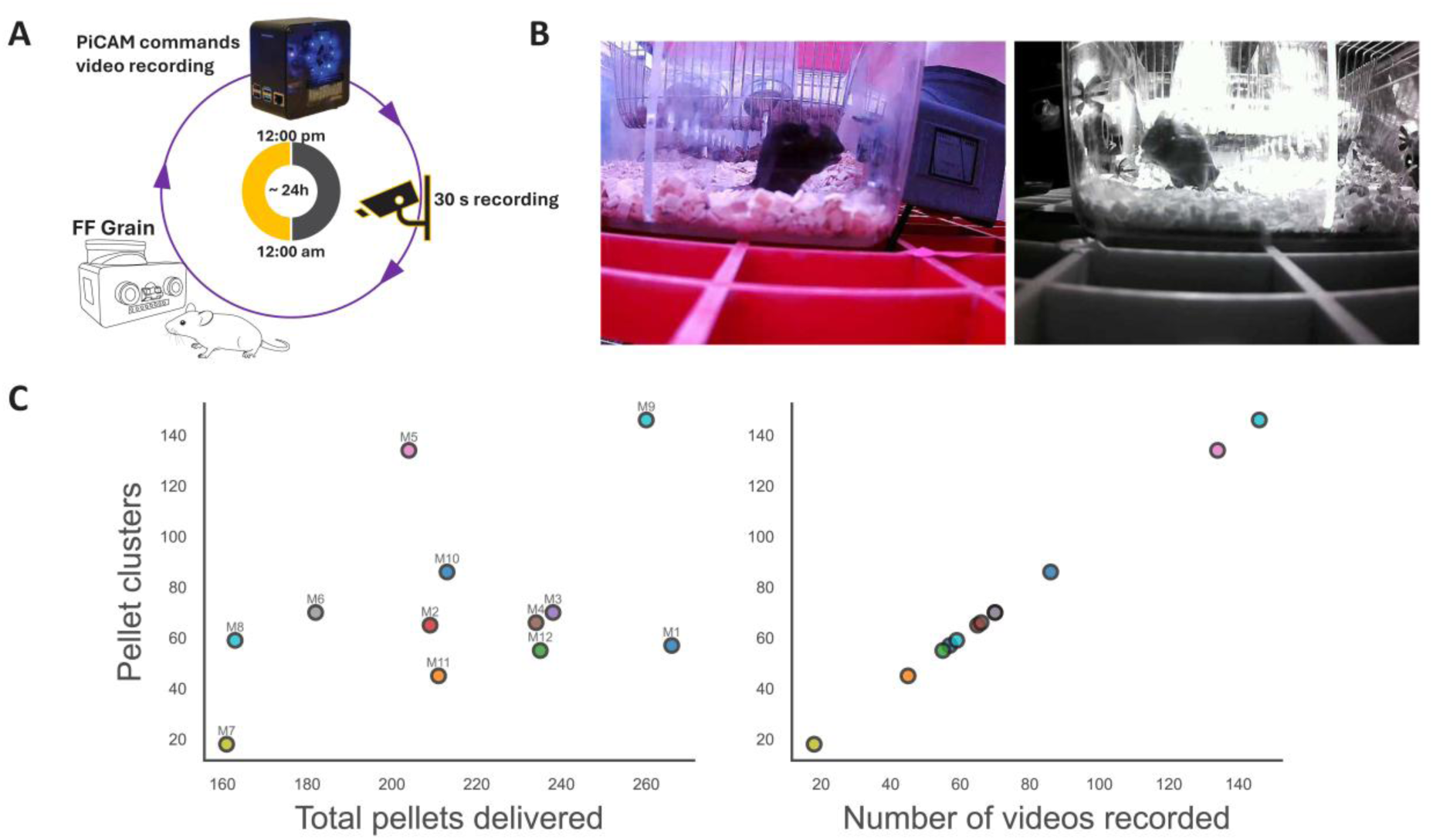
Validation of the RTFED (PiCAM) system. (**A**) Schematic representation of the setup, each mouse had access to a FED3 filled with 20 mg grain pellets on FF mode for ∼24 hours and every pellet retrieval was recorded using PiCAM recording module. (**B**) Snapshots of pellet consumption events captured at day (Left) and night (Right) by RTFED (PiCAM). (**C**) The relationship between total number of pellets collected and number of pellet clusters (triggered events) per mouse (Left panel). Each point represents one animal (M1–M12). Labels correspond to mouse IDs. Video recording was triggered by pellet retrieval events and extended if additional pellets were collected within 30 seconds. The number of clusters corresponded exactly to the number of videos saved in the experimental folders (Right panel), confirming precise video– event alignment.

### 3.3 Experiment 3: Studying association between behavioural events and dopamine release using RTFED (PiTTL)

#### 3.3.1 Benchmarking of TTLs in an artificial setting

Before running *Experiment 3*, we evaluated the precision and reliability of TTL signal transmission with RTFED (PiTTL). To do this, we systematically tested event-triggered TTL delivery in response to mimicked interactions on four FED3s connected to the Raspberry Pi. Latency measurements were implemented directly within the TTL transmission functions of RTFED (PiTTL) original GUI code (Fig. 7**A**). Briefly, the timing of each event trigger and corresponding TTL pulse was logged with high precision. TTLs were generated on different GPIO pins assigned to individual FED3 ports. On each FED3 unit, we simulated more than 20 pokes and pellet delivery events in different combinations and timing, separately or simultaneously. The output of the test was saved as .csv file and to visualize system responsiveness, we plotted a kernel density estimate (KDE) of event-to-TTL delays (Fig. 7**B**).

**Fig. 7.**
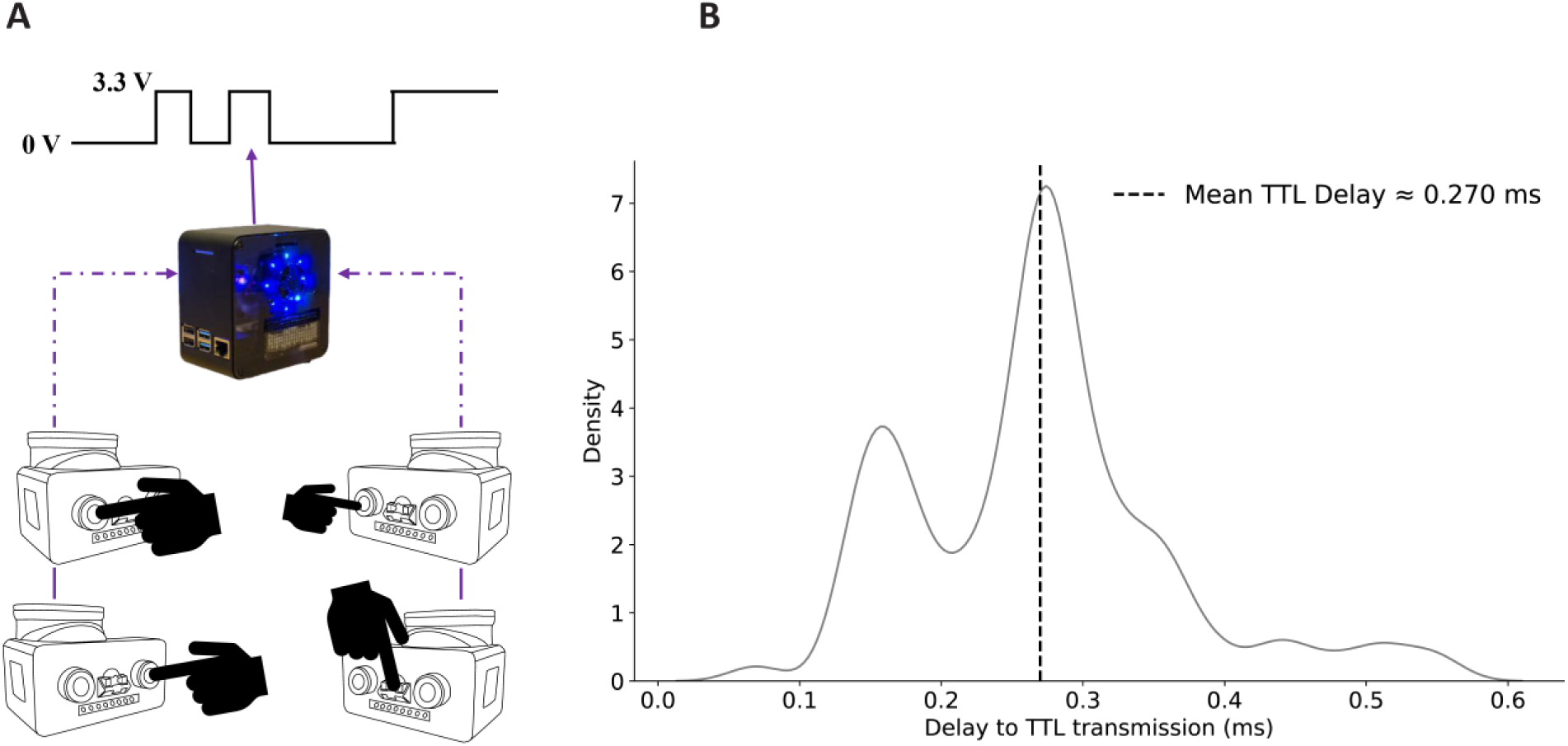
Benchmarking the TTL transmission rate. (A) Schematic representation of the experiment, ∼100 interactions were mimicked on 4 FED3 devices connected to the RTFED (PiTTL) via USB cables and the latency to activate pins were logged. (B) A KDE plot of all events shows that the mean latency to transmit the TTL signal from FED3s to the pins on the Raspberry Pi is 0.27 ms.

The mean event-to-TTL delay across all ports was 0.270 ms. The KDE plot showed that the majority of TTLs were sent within 0.3 seconds with minimal jitter or port-based variability (Table. S1). No transmission failures or timing outliers were observed in any trial. This confirms that the PiTTL system provides sub-millisecond latency, more than sufficient for precise alignment with fibre photometry or other closed-loop setups.

#### 3.3.2 Measurement of dopaminergic activity in NAc of mice using RTFED (PiTTL)

In the final experiment (Fig. 8**A**), mice expressing *GRAB_DA_* in NAc core were used to test the PiTTL module of RTFED. Recordings were carried out in the home-cage of mice and the mice were free to explore and interact with FED3s (Supplementary video V5-6). The PiTTL module successfully delivered TTL pulses across several sessions and the group-level analysis showed difference in dopamine release in the NAc core in response to grain and sucrose pellets (**Fig. 8B-C**).

**Fig. 8.**
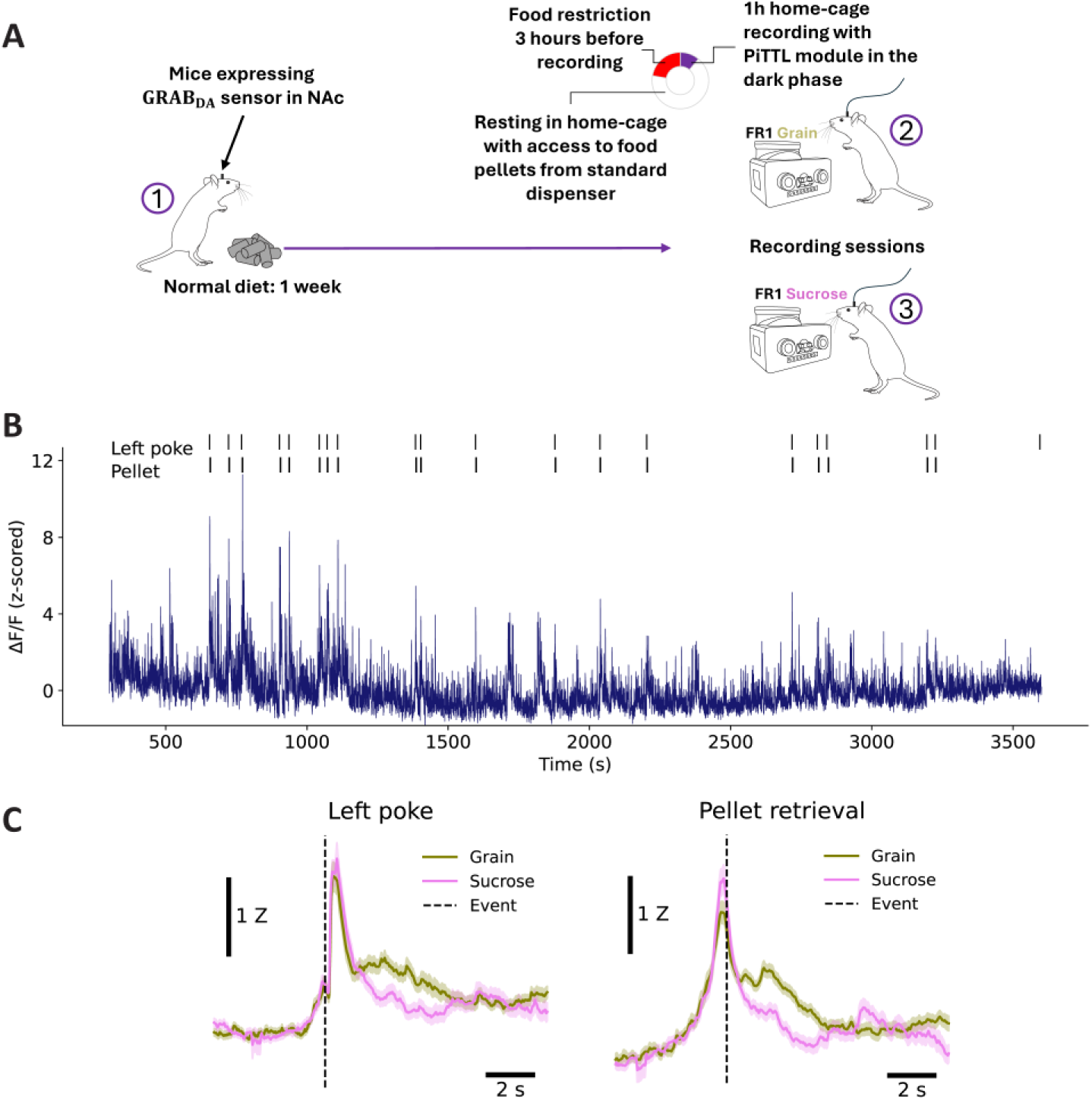
PiTTL integration with photometry setup. (A) Study paradigm of *Experiment 3*. After recovery from surgery, mice were first maintained on normal diet with ad lib access to food and water for 1 week. 3 hours before recording session, food pellets were removed from the dispenser and mice were moved to the recording room, FED3s were switched on and connected to the PiTTL system. Mice could make a left poke to earn a grain pellet. One day later, another recording session with sucrose pellets was carried out. (B) A representative raw trace of a full recording session of a single mouse with an overlay of TTL pulses received from PiTTL during the session. (C) Traces were averaged across trials of all mice and aligned to Left poke and the Pellet retrieval events.

## 4. Discussion

The transition toward advanced automated behavioural analysis underscores the necessity for robust, continuous, and unobtrusive data collection approaches. For example, decision-making studies, such as those involving probabilistic reward schedules, benefit from longitudinal data collection without human interference (St Onge & Floresco, 2009) or various factors that affects metabolism or feeding behaviour cannot be studied without the use of home-cage monitoring system (Tschöp et al., 2011). Longitudinal observations enabled by these home-cage systems have proven invaluable, offering stable and individualized assessments that inform vulnerabilities or resilience patterns in response to various environmental or physiological stressors (Jhuang et al., 2010; Kahnau et al., 2023).

In this study, we presented RTFED, a novel open-source enhancement for the widely used FED3 device, aimed at significantly advancing real-time and remote behavioural monitoring capabilities in mice. By addressing critical limitations inherent to traditional methodologies (e.g., manual weighing of food, continuous video recording, and use of experimental chambers) and the original FED3 firmware, RTFED ensures continuous, robust, and stress-minimized data acquisition, crucial for investigating naturalistic feeding patterns and operant behaviours within home-cage environments. Here we focused on the most common feeding-related tasks e.g., pellet retrieval and pre-programmed operant tasks, however, theoretically use of RTFED could be expanded to cover any program running on the FED3, for example, social behaviour studies as described in Chen and colleagues who used FED3 to design a social operant task to study social motivation for multiple days in the home-cage (Chen et al., 2025).

Our validation experiments demonstrated the reliability, precision, and versatility of the RTFED system across various configurations. The real-time logging feature seamlessly integrated with cloud-based Google Sheets, enabling automated and remote monitoring of behavioural events without manual intervention, thus facilitating large-scale studies with more innovative study paradigms. Nevertheless, it is worth noting that, RTFED is designed to work with Google products, thus it can be a limitation for labs whose institutions prohibit using such services due to their own IT restrictions. Moreover, the RTFED (Basic) is built on the architecture of Windows environment, which means that users running Mac systems cannot benefit from it.

The RTFED (PiCAM) configuration, equipped with event-triggered video recording, precisely captured pellet retrieval events, and the numbers of videos perfectly matched expected clusters derived from feeding logs, demonstrating high accuracy and robust synchronization between behavioural events and recorded footage. Furthermore, the RTFED (PiTTL) implementation provided precise and sub-millisecond accurate TTL pulse transmission, crucial for integration with advanced neurophysiological methods such as fibre photometry and closed-loop protocols. Benchmarking confirmed minimal jitter, negligible latency variability, and high reliability, validating its application in real-time neuroscience paradigms.

Importantly, RTFED aligns with the broader shift in neuroscience from reliance on costly, closed-source commercial platforms toward adaptable, open-source technologies. Recent trends illustrate that Raspberry Pi-based and other DIY automated solutions have enabled comprehensive measurements including body weight, fluid consumption, and detailed behavioural tracking with remarkable flexibility and affordability (Godynyuk et al., 2019; London et al., 2018). Moreover, incorporating artificial intelligence into home-cage monitoring setups has allowed for the integration of complex behavioural analytics and neurophysiological recordings, providing deeper insights into the dynamic interplay between behaviour and brain activity (Pereira et al., 2022; Spangenberg & Keeling, 2016).

Overall, RTFED offers a powerful yet user-friendly solution to contemporary behavioural neuroscience challenges, bridging gaps left by commercial and existing open-source tools. Its ease of use, low-cost requirements, adaptability, and robust feature set are expected to significantly broaden accessibility, fostering innovation, and accelerating discoveries in behavioural neuroscience research.

In summary, the evolution of automated home-cage behavioural monitoring tools, represents a significant advancement in neuroscience. While FED3 has provided foundational improvements, ongoing limitations necessitate continued innovation. Future developments should focus on expanding integrative capabilities, particularly leveraging machine learning and wireless sensor technologies. For example, freely available machine-learning tools can be used to train pose-estimation models and behavioural classifiers integrated into RTFED (PiCAM + PiTTL) enhanced with GPU-powered Jetson boards to detect particular movements of mice such as extending the paw to collect the pellets or other post-consummatory behaviours like grooming, drinking or sleeping, and transmit those behaviours as TTL pulses to the neural recording units. This will promote further understanding of complex behaviour-brain relationships with translational relevance.

## Acknowledgments

The authors would like to acknowledge Eloise Kuijer for user-experience feedback that helped us tailor the GUI for a better performance, and Tobias Alexander Nikolaisen for modifying the 3D model of the camera mount and initial benchmarking of the PiCAM system and feedback along with Karin Linnea Volcko who ran additional benchmarking of the PiCAM system. We would like to thank Alexxai V. Kravitz as the original author of FED3 as well as Fabrice de Chaumont for initial discussions about real-time data collection from FED3 units. Moreover, we would like to thank the TEATIME action of COST for supporting and encouraging researchers to develop home-cage monitoring systems. We also wish to thank the staff at the animal facility of UiT and workshops of the Faculty of Health Sciences at UiT.

