## Supplementary File Text for "RTFED, an open-source versatile tool for home-cage monitoring of behaviour and fibre photometry recording in mice"

The materials in this document, including code snippets, figures, videos and tables are arranged in the order of their first appearance in the main manuscript.

### 2.3.S1. Serial communication initialization and sleep management

Here, we initially cover all the major changes made in the firmware, and then we explain each adjustment with more details in the following paragraphs. Firstly, to ensure continuous connectivity and support real-time data exchange between FED3 and a host computer or a microprocessor, we disabled the default FED3 sleep behaviour by adding a disableSleep() call in the FED3::begin() method and increased the serial baud rate from 9600 to 115200. These changes allow the device to remain actively connected to a host Windows computer or Raspberry Pi over serial communication, which is critical for transmitting behavioural events as they occur and is the foundation for online data logging and remote monitoring of behaviour (which ultimately supports TTL pulse generation, and event-triggered video recording of behaviours). An important note is that users should not call the sleep() function in their customized scripts, otherwise FEDs will be disconnected from the RTFED software. Secondly, we introduced a set of new firmware features that enhance both experimental control and system reliability. The new RTFED library enables FED3 units to listen to serial commands (e.g., for clock synchronization across multiple devices or easy mode selection) to ensure precise timing across multiple FED3s or switching modes by one master command to all FED3s. The clock synchronization feature is not only useful for proper data management, but also highly important for experiments involving timed feeding (start and stop feeding at a particular hour) or choice paradigms where the animal collects reward from multiple devices and the choice patterns cannot be interpreted if devices are not synchronized to each other. We also added new event types, including "PelletInWell" to mark the onset and offset of pellet delivery to the well and "JAM" to detect dispensing failures. In case of a jamming event, which basically disrupts data collection, the new firmware logs the error, halts further dispensing attempts, and triggers an alarm, which can be configured to send an email notification to alert the user. These additional events and controls allow researchers to continuously monitor and respond to feeding behaviours and device status in real time, making RTFED more robust than the original FED3 firmware.

The original FED3 only remains connected to a computer if put on bootloader mode, which is primarily used to flash the board (e.g., installing new programs). Otherwise, it goes to sleep every 5 seconds to save battery life. In our RTFED system, the FED3s are always plugged via USB cables to either a Windows computer or a Raspberry Pi, so sustaining battery life is irrelevant. More importantly, sleep mode shuts down serial communication. Therefore, we decided to disable sleep mode entirely.

In the original FED3 library:

```
void FED3::begin() {
   Serial.begin(9600); // Uses a slower baud rate
   // No explicit call to disable sleep, so the device may sleep and
      disconnect the USB serial link
}
```

While in the RTFED library:

```
void FED3::begin() {
  Serial.begin(115200); // Higher baud rate for faster, more reliable
     serial communication
  disableSleep();       // Ensures the device remains awake (keeping
     the USB connection active)
}
```

RTFED uses a baud rate of 115200 compared to 9600 in the original FED3, improving speed and reliability, especially for TTL transmission. Calling `disableSleep()` ensures the device stays connected for real-time monitoring. Users must avoid calling the sleep function in custom scripts, or else the FED will disconnect and no longer be recognised by RTFED.

### 2.3.S2. Enhancing data logging via serial output

In the original FED3, users only accessed data via the SD card after the experiment. In contrast, RTFED enables real-time serial logging for immediate observation and debugging:

```
void FED3::logdata() {
  // ... data preparation ...
  String logData = /* formatted log string with date, event type,
     counts, etc. */;
  Serial.println(logData);       // Outputs to serial monitor
  logfile.println(logData);      // Logs to SD card
}
```

This allows real-time feedback while preserving backup copies on both SD card and local computer.

### 2.3.S3. Listening for serial commands (Time synchronization, Simulated pokes and Mode selection)

FED3 has a coin battery-powered clock for timestamping, but it can drift or reset, causing issues for synchronisation across multiple devices. RTFED defines a function to process serial commands and solve this problem.

`checkSerialCommands` handles:

- `SET_TIME`: set real-time clock (RTC)
- `SET_MODE`: set FED3 mode (0–12)
- `TRIGGER_POKE`: simulate left or right poke

```
void FED3::checkSerialCommands() {
  if (Serial.available()) {
    String line = Serial.readStringUntil('\n');
    line.trim();

    if (line.startsWith("SET_TIME:")) {
      line.remove(0, 9);
      int year, month, day, hour, minute, second;
      char cstr[50];
      line.toCharArray(cstr, sizeof(cstr));
      int parsed = sscanf(cstr, "%d,%d,%d,%d,%d,%d",
                          &year, &month, &day,
                          &hour, &minute, &second);
      if (parsed == 6) {
        rtc.adjust(DateTime(year, month, day, hour, minute, second));
        Serial.println("TIME_SET_OK");
      } else {
        Serial.println("TIME_SET_FAIL");
      }

    } else if (line.startsWith("SET_MODE:")) {
```

```
      line.remove(0, 9);
      int mode = line.toInt();
      if (mode >= 0 && mode <= 12) {
        FEDmode = mode;
        writeFEDmode();
        Serial.println("MODE_SET_OK");
        delay(200);
        NVIC_SystemReset();
      } else {
        Serial.println("MODE_SET_FAIL");
      }

    } else if (line == "TRIGGER_POKE") {
      simulatePoke();
    }
  }
}
```

```
void FED3::run() {
  if (stopLogging){
    return;
  }
  checkSerialCommands();
}
```

The `run()` function integrates `checkSerialCommands()` into the device's main loop, enabling live control and configuration of experiments.

### 2.3.S4. Additional events defined for better monitoring of behaviour and handling of failures

RTFED introduces two new event types:

- `PelletInWell`: when a pellet is detected in the well
- `JAM`: when pellet dispensing fails

```
if (pelletDispensed == true) {
  ReleaseMotor();
  pelletTime = millis();
  Event = "PelletInWell";
  logdata();
}
```

If delivery fails, a series of recovery steps is attempted, and a JAM event is triggered after repeated failures:

```
if (PelletAvailable == false) {
  pelletDispensed = dispenseTimer_ms(1500);
  numMotorTurns++;

  if (pelletDispensed == false) {
    if (numMotorTurns % 5 == 0) pelletDispensed = MinorJam();
    if (numMotorTurns % 10 == 0) pelletDispensed = VibrateJam();
    if (numMotorTurns % 20 == 0) pelletDispensed = ClearJam();
```

```
    if (numMotorTurns % 50 == 0) {
      Alarm();
    }
  }
}
```

`Alarm()` logs the event, halts further actions, and disables the system:

```
void FED3::Alarm() {
  Event = "JAM";
  logdata();
  stopLogging = true;
  disableInputs();
  jamOccurred = true;
}
```

Unlike the original firmware which looped silently during jams, RTFED logs the error and can notify users via email.

### 2.3.S5. Extending modes for RTFED and GUI integration

If you need to have more modes included in your library and then recognized by the RTFED GUI, adjust these snippets:

```
// SET_MODE:<0-15>
} else if (line.startsWith("SET_MODE:")) {
  line.remove(0, 9); // remove "SET_MODE:"
  int mode = line.toInt();
  if (mode >= 0 && mode <= 15) {
    FEDmode = mode;
    writeFEDmode();              // Save new mode to SD
    Serial.println("MODE_SET_OK");
    delay(200);
    NVIC_SystemReset();          // Reboot into new mode
  } else {
    Serial.println("MODE_SET_FAIL");
  }
}
```

Here you can extend the valid range for modes 16, 17, etc.

```
// Display selected mode
display.setCursor(10, 60);
if (FEDmode == 0)  display.print("Mode 1");
if (FEDmode == 1)  display.print("Mode 2");
% ... up through ...
if (FEDmode == 15) display.print("Mode 16");
// ...and so on for any extra modes...
DisplayMouse();
display.clearDisplay();
display.refresh();
```

```
// Classic mode labels
if (ClassicFED3 == true) {
  if (FEDmode == 0)  display.print("Free feeding");
  if (FEDmode == 1)  display.print("FR1");
  % ... up through ...
  if (FEDmode == 15) display.print("DetBandit");
```

```
  // ...add extra labels here...
  display.refresh();
}
```

```
// Set FR based on FEDmode
if (FEDmode == 0)  FR = 0;    // free feeding
if (FEDmode == 1)  FR = 1;    // FR1
% ... up through ...
if (FEDmode >= 6 && FEDmode <= 15) FR = 1;
// ...for new modes, set FR accordingly...
```

Finally, in the RTFED GUI (Python), extend the mode list:

```
self.mode_options = [
    "0 - Free Feeding", "1 - FR1", "2 - FR3", "3 - FR5",
    "4 - Progressive Ratio", "5 - Extinction", "6 - Light Tracking",
    "7 - FR1 (Reversed)", "8 - PR (Reversed)", "9 - Self-Stim",
    "10 - Self-Stim (Reversed)", "11 - Timed Feeding",
    "12 - ClosedEconomy_PR2", "13 - Probabilistic Reversal",
    "14 - Bandit8020", "15 - DetBandit",
    # ...add "16 - YourNewMode", "17 - AnotherMode", etc.
]
```

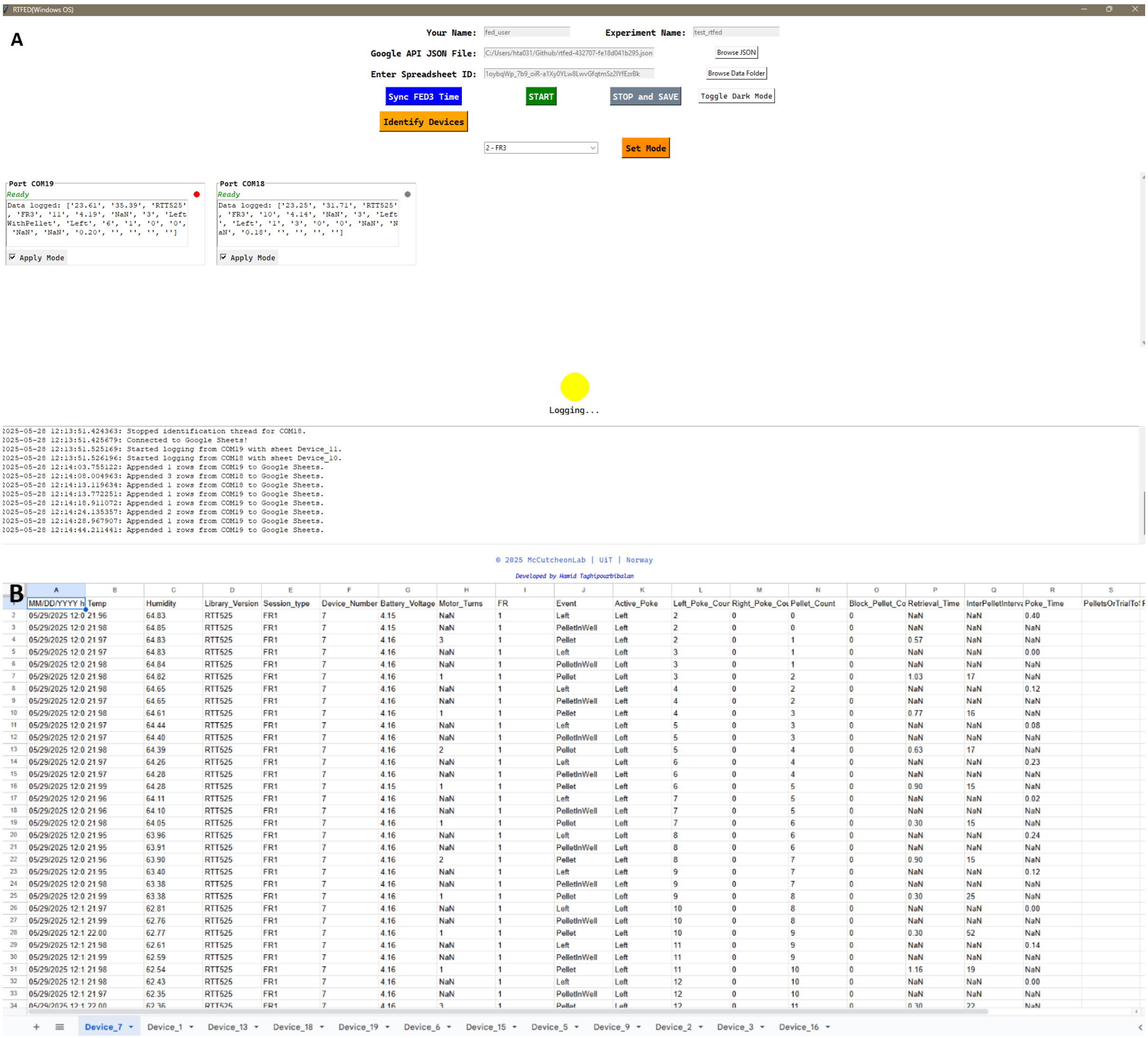

Fig. S1. (A) The RTFED GUI offers a user-friendly set of tools which makes the data management of experiments much easier, allows synchronization of the time on all FED3 units, and facilities the process of setting new modes on FED3 units by pressing a button instead of making separate pokes on all devices. (B) The RTFED GUI also logs the data on Google Sheets labelled with device number instead of separate CSV files, nevertheless the data is also saved as separate CSV files on the local machine in addition to each FED3s' microSD card. Upon launching the RTFED(Basic) GUI the users begin by entering their name and experiment title, selecting a Google API JSON credential file and inputting the Google Spreadsheet ID where data will be stored online. The GUI automatically detects connected FED3 units via USB and displays them in scrollable frames, each showing real-time status, event logs, and activity indicators. A dedicated "Identify Devices" button triggers a poke signal on each FED3, enabling the system to associate each port with its unique Device Number before logging begins. The "Sync FED3 Time" function allows users to synchronise all devices clocks with the host computer. Additionally, using a drop-down menu, the user can select their desired mode for either all or a selected group of FED3 units which would restart and set the new modes on the selected FED3s. Pressing "START" initiates real-time data logging to Google Sheets, with device-specific worksheets created automatically. A circular indicator shows the current recording status (standby, logging, or off), and a live log viewer tracks system messages. On stopping the session with "STOP and SAVE" button, all the data will also be saved locally, and serial connections will be safely closed.

Video. V1. A tutorial demonstration of RTFED(BASIC) GUI on Windows OS. The video shows how to setup an experiment on the GUI, synchronize FED3 devices and utilize the other features of the GUI.

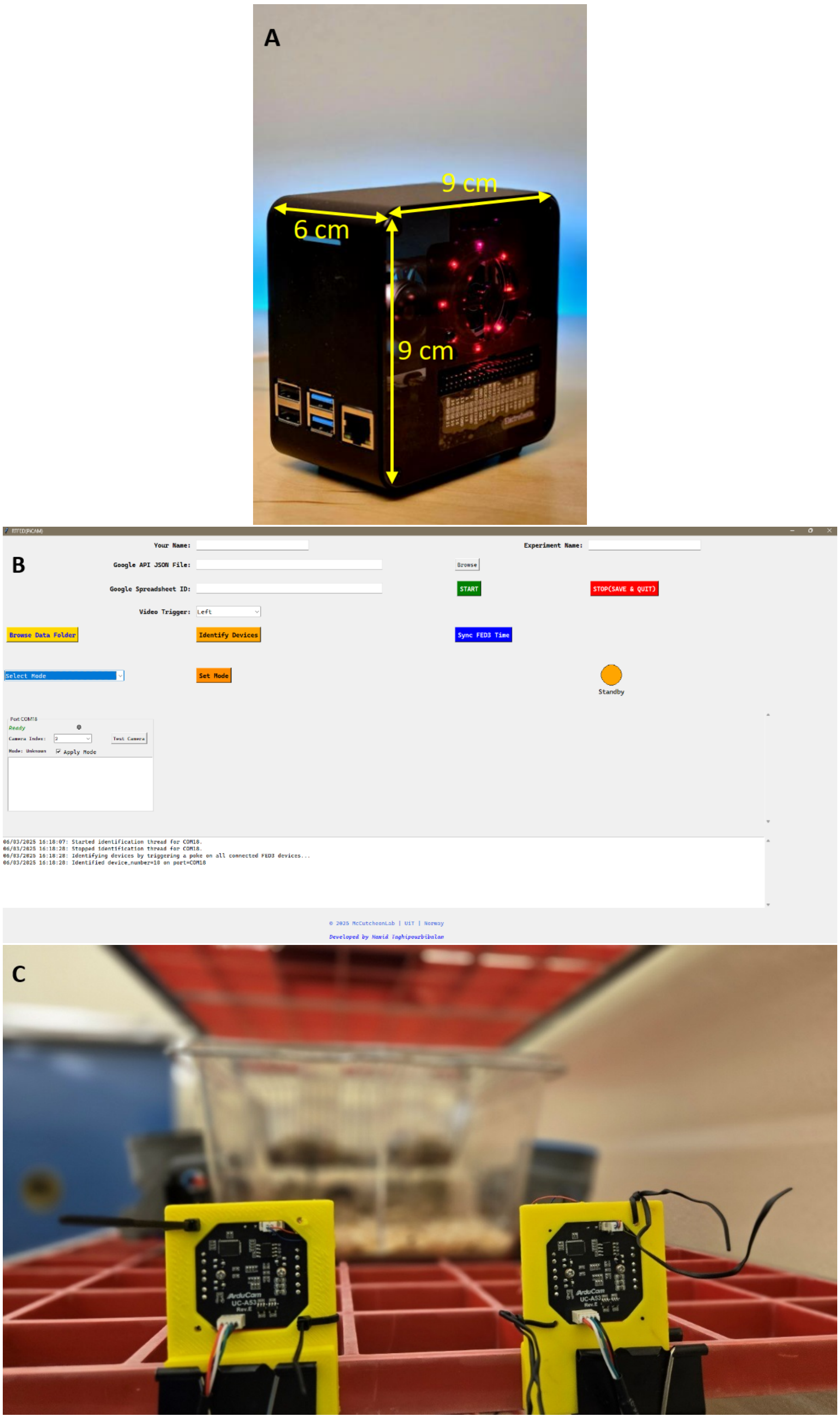


Fig. S2. (A) The Raspberry Pi 4B is a compact single board computer that can be purchased together with a case and cooling system for a very cost-efficient price. (B)The RTFED (Pi-CAM) GUI offers an additional feature to the RTFED(Basic) which is the ability to initiate event-triggered video recording of behaviours such as nose pokes and pellet retrieval. (C)The user can plug USB cameras along with the FEDs to the Raspberry Pi and couple the cameras with each FED3 unit.

Video. V2. The PiTTL system works as a relay station to transmit TTL pulses from FED3 devices,this simple benchmarking shows how the interactions with FED3 devices are transmitted via the GPIOs of the Raspberry Pi, displayed via the output LEDs.

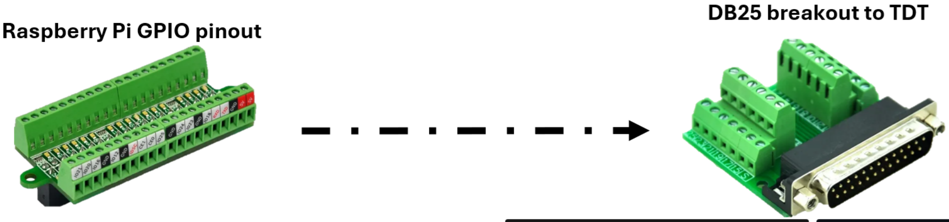


```
gpio_pins_per_device = {
    'Port 1': {"LeftPoke": 17, "RightPoke": 27, "Pellet": 22},
    'Port 2': {"LeftPoke": 10, "RightPoke": 9,  "Pellet": 11},
    'Port 3': {"LeftPoke": 0,  "RightPoke": 5,  "Pellet": 6},
    'Port 4': {"LeftPoke": 13, "RightPoke": 19, "Pellet": 26},
    'Port 5': {"LeftPoke": 14, "RightPoke": 15, "Pellet": 18},
    'Port 6': {"LeftPoke": 23, "RightPoke": 24, "Pellet": 25},
    'Port 7': {"LeftPoke": 8,  "RightPoke": 7,  "Pellet": 1},
    'Port 8': {"LeftPoke": 12, "RightPoke": 16, "Pellet": 20},
}
```

Port 1/2

| DB25 Pin | Pi pin |
|---|---|
| 1 | Right 27 |
| 14 | Left 17 |
| 2 | Pellet 22 |
| 15 | Right 9 |
| 3 | Left 10 |
| 16 | Pellet 11 |
| 5 | Ground |

Port 3/4

| DB25 Pin | Pi pin |
|---|---|
| 1 | Right 5 |
| 14 | Left 0 |
| 2 | Pellet 6 |
| 15 | Right 19 |
| 3 | Left 13 |
| 16 | Pellet 26 |
| 5 | Ground |

Fig. S3. The map of connections from Raspberry Pi 4B pins to the DB25 breakout terminal. We recommend using the ultra-small RPi GPIO Status LED and Terminal block breakout board listed in Table 1. This module enables the users to easily and securely connect the RPi pins to the DB25 breakout using jumper wires. Additionally, the module comes with status LEDs, which are ideal for ensuring the TTLs are being transmitted via the correct pin.

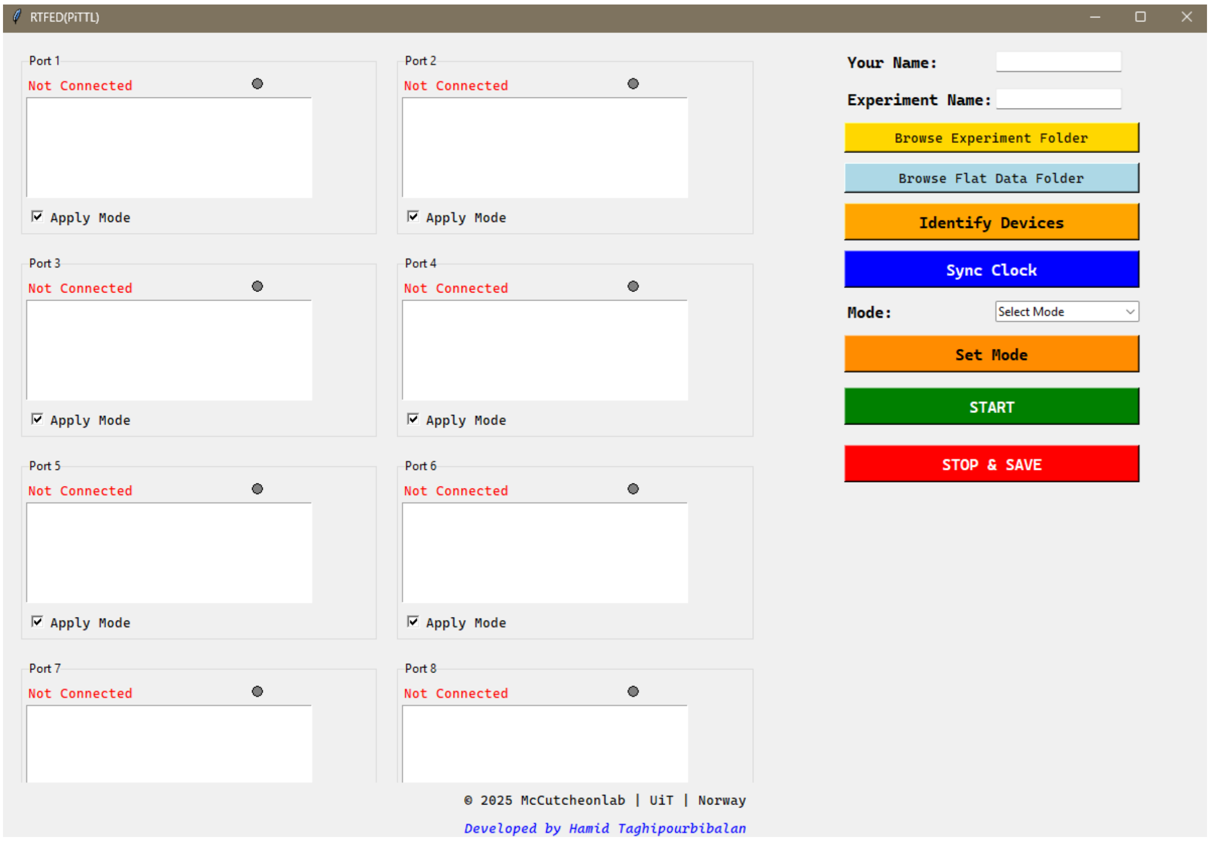


Fig. S4. The RTFED (PiTTL) GUI offers most of the basic features of RTFED system, handles TTL pulses to the photometry system, and, for long-term recordings, it also allows setting up online data collection. However, this feature is by default disabled on the GUI and the user can toggle it on.

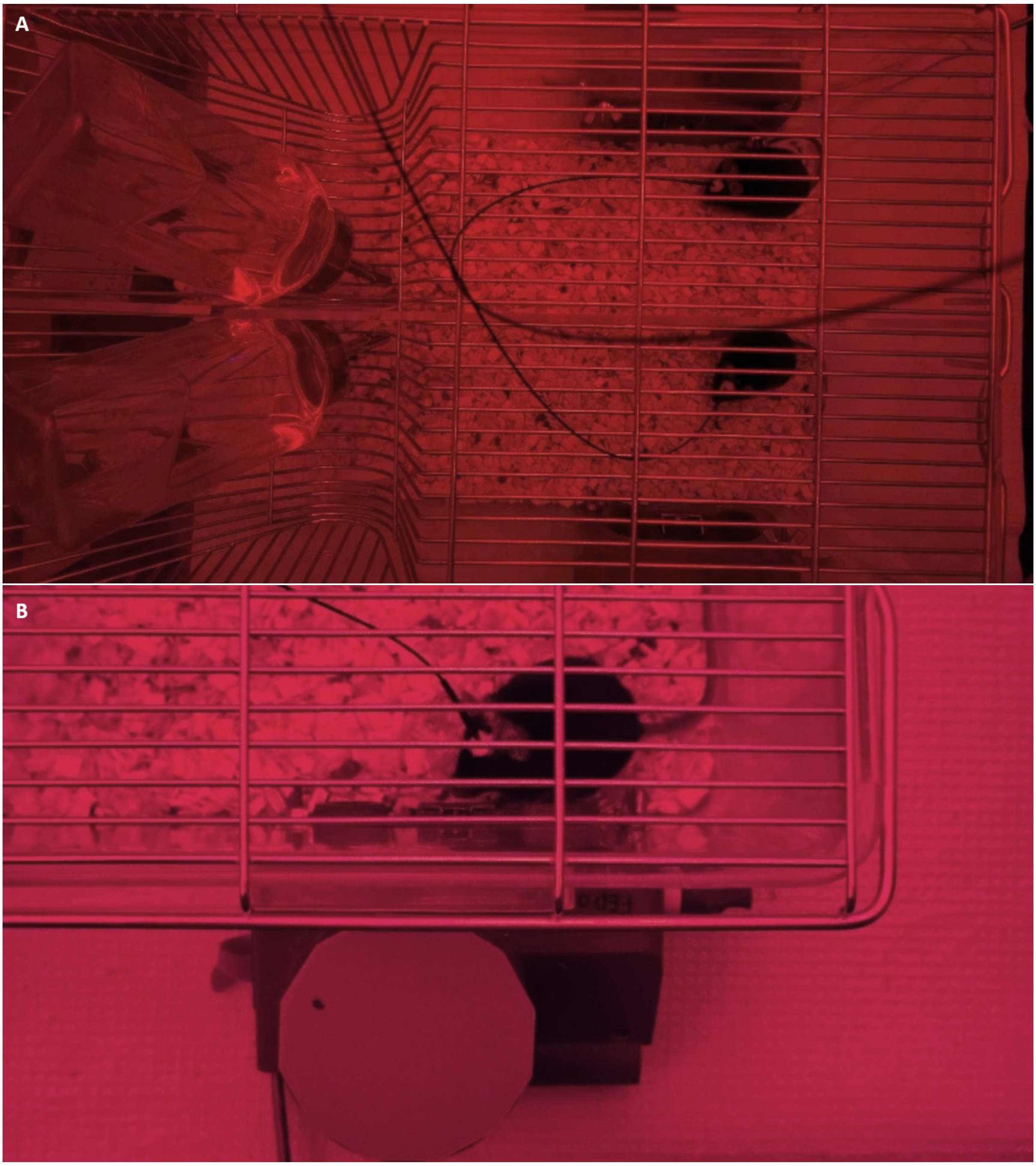


Fig. S5. (A) Two mice per cage separated by a divider were recorded at the same time each with access to their own FED3. (B) A mouse plugged into the photometry system collects pellets from FED3 while being recorded using the PiTTL system.

Video. V3. The video shows two mice in a contact-housed setting plugged into the fibre photometry system and being recorded using FED3 and PiTTL.

Video. V4. Fast-forwarded video of a mouse collecting multiple pellets recorded in a single file by PiCAM.

Table. S1. The table shows the mean of the TTL delay across the 4 tested ports on a Raspberry Pi 4 B.

| Port # | Mean delay | SD delay | Mean duration | SD duration | N |
|---|---|---|---|---|---|
| **Port 1** | 0.000242391 | 8.638220520161587e-05 | 0.10019221739130435 | 9.39206418021148e-05 | 23 |
| **Port 2** | 0.00027892 | 9.484677116275496e-05 | 0.10022492 | 0.000196539 | 25 |
| **Port 3** | 0.000308222 | 0.000109588 | 0.10018440740740742 | 9.244510190429971e-05 | 27 |
| **Port 4** | 0.000249692 | 6.831735898336188e-05 | 0.10019253846153846 | 0.000102958 | 26 |

Video. V5. Mice can freely explore the cage, drink water, rest in the nesting material and interact with FED3s on their own while they are plugged into the recording system.

Video. V6. The overlay video of a photometry signal aligned with the behaviour using PiTTL system. First the mouse makes a left poke which is marked by auditory (beep sound) and visual(light) cues and reflected as TTL pulse, right after the poke, the mouse collects a pellet which follows a surge in the dopamine signal(blue signal).
